# A novel bioactive molecule 5-β-Hydroxyltridecanyl benzoate identified in Pedalium Murex L. (Gokhru) fruit extracts showed antifungal activity against *Sclerotinia sclerotiorum*

**DOI:** 10.1101/381244

**Authors:** Gulshan Chaudhary, Nutan Kaushik, Pallavolu Maheswara Reddy

## Abstract

Fruits of *Pedalium murex* known as, ‘Gokhru’ is predominantly used in Ayurveda for its aphrodisiac properties and use in treatment of urological orders. However till now, bioactivity of this fruit has not been documented for plant protection against phytopathogenes, Hence in this regard, we investigated antimicrobial activity of *P. murex* fruit extracts (isolated from the different agro climatic zones of India) were tested against twelve microbial phytopathogens. From 29 fruit extracts, only 13 were found to be active against three pathogens viz. *Botrytis cinerea, Rhizoctonia solani*, and *Sclerotinia sclerotiorum*. Among the nine TLC fractions only one showed presence of a single peak, thus presence of a single compound in the crude extract. Based on lower IC_50_ value (328 μg/ml) the crude fruit extract collected from Indore region was further characterized. Evaluation against screened phytopathogens indicates its activity. Further, structural characterization of this compound using FTIR and NMR revealed it to be a novel compound i.e. 5-β-Hydroxyl tridecanyl benzoate. Our study paves a way to investigate effective dose of *P. murex* fruit extracts to utilize in plant protection programs. In future, a better method is required to scale up the process of bioactive molecule (5-β-Hydroxyl tridecayl benzoate) extraction from *P. murex* or for its chemical synthesis.

## Introduction

Plants are known for containment of bioactive metabolites which contributes to plant defense mechanisms against pests. Most of the metabolites give plants their odors (terpenoids), some are responsible for plant pigments (quinines and tannins), and others (e.g., some of the terpenoids) are responsible for plant flavor. These bioactive compounds or secondary plant metabolites consist of low-molecular-weight compounds that are regarded as not essential for sustaining life, but as crucial for the survival of the organism (Hadacek 2002). More than 50,000 structures have been identified in plants by various chemical analysis. However, as only less than 20% of all plants have been studied, it is very likely that the actual numbers of secondary metabolites or bioactive compounds in the plant kingdom would exceed 100,000 structures (Wink 2006). Among these, the antimicrobial type of bioactive compounds is commonly divided into five main classes consisting: terpenoids and essential oils; phenolics and polyphenols; alkaloids; polypeptides and mixtures (crude extract) (Zacchino et al. 2017).

Plant pathogenic fungi cause severe crop damages both at early developmental stages to post-harvest stages. A range of fungal genera is responsible for the compromise in crop production and other aspects including nutritional value, organoleptic characteristics, and reduced shelf life (Agrios 2004). Simultaneously, fungi are also indirectly responsible for allergic or toxic disorders among consumers because of molecules production which is characterized as allergens or mycotoxins (Harris et al. 2001). In the past, plant pathogens are handled with a variety of synthetic fungicides to improve agricultural output, but benefits always come with side effects in the form of carcinogenicity, residual toxicities and other detrimental effects on the environment. Thus alternative and non-hazardous choice for the use of plant-based natural fungicides would be advantageous, which are non-phytotoxic, systematic and readily biodegradable. Hence, there is a high demand for novel antifungals belonging to a wide range of structural classes, selectively acting on new targets with fewer side effects (Abad et al. 2007). According to current estimates, about 10 to 20% of staple foods and cash crops are destroyed by plant pathogens (Pratiwi et al. 2018). Currently, there is little evidence on the antimicrobial properties of the medicinal plants under investigation against phytopathogen fungi.

Among several pathogenic fungi, *Rhizoctonia solani, Sclerotinia sclerotiorum* and *Botrytis cinerea* are major plant pathogens. *Rhizoctonia solani* causes significant establishment and yield losses to several essential food crops globally (Wibberg et al. 2016). Necrotrophic fungal pathogen *Sclerotinias clerotiorum* (Lib.) is one of the most devastating and cosmopolitan soil borne Ascomycetes, infecting over 500 species of plants worldwide. Annual yield losses due to *Sclerotinia* diseases exceed over several hundred million dollars each year world over (Zhu et al. 2016). Fungicides currently in use are unable to provide complete disease control on sclerotia of *S. sclerotiorum* due to the lack of ability to move systematically in plants (Peltier, 2012). Another necrotrophic fungus *B. cinerea* is well known for its extensive host range, wide distribution globally, extreme variability and adaptability to extensive environmental conditions (Clarkson et al., 2017). *Botrytis cinerea* is the causal agent of gray mold disease in a broad range of dicotyledonous plants (Rupp et al. 2017). It colonizes senescent or wounded tissues but is also able to infect healthy plants, causing serious damage in fruits and vegetables in open fields and in greenhouses, both during pre- and post-harvest (Droby and Licter 2007). *B. cinerea* is considered to be an exemplary necrotroph, as it meets all the classic criteria of this type of pathogen (Tudzyski and Kokkelink 2009). Its infection strategy includes the killing of host cells using the secretion of cell wall degrading enzymes and toxic metabolites that induce cell death in advance of the invading hyphae (Kars and Van Kan 2007; Benito-pescador et al. 2016).

*Pedalium murex* L, commonly known as Bada Gokhru, belongs to Pedaliaceae family and is known for its pharmacological uses (aphrodisiac and in urological disorders)in traditional medicine system (Chaudhary and Kaushik 2017). The plant is very well known for pharmacological research. However, less focus is given on its activity as bio-pesticide although the plant is reported for its insecticidal activities (Sahayarajet al, 2008; Raja et al, 2007). The plant is reported to have antifungal activity against human pathogens (Rajashekar et al. 2012; Muruganatham 2011). However, some reports are also available for its activity against plant pathogens such as *Magnaporthe grisea* for its antifungal properties (Parimelazhagan 2011). Therefore, the present study focused on assessment of antifungal and antibacterial properties of the extract prepared from fruits of *P. murex* collected from diverse locations of India and aim to isolate and to characterize the bioactive compound with chemical analysis.

## Materials and methods

### The material used in the study

Twenty-nine fruit samples of *P. murex* were collected from different agro climatic locations from ten states of India (Table S1). All collected fruits were submitted to National Institute of Science Communication and Information Resources (NISCAIR), New Delhi for the authentication and use for further investigations. Various Plant pathogens (fungi and bacteria) of agro-economic importance were collected from Indian Type Culture Collection (ITCC), Indian Agricultural Research Institute (IARI) (Table S2) and were maintained on Potato Dextrose Agar (PDA) and Nutrient Agar Medium (NAM) as described by Haynes et al. (1995).

### Preparation of plant extracts

Approximately, 50-100 gram of mature *P. murex* fruits were washed with water, shade dried for 2-3 days, and powdered in ultra centrifugal mill ZM 200, (Retsch, Germany). The fruit powders were extracted in methanol (~300-700ml; varies according to the amount of fruit powder) in Soxhlet apparatus for 48 h and concentrated using rotary evaporator (Heidolph, Germany). The extracts were stored at 4°C till further analysis.

### Screening of extract for antimicrobial activity

Bioassays were performed to test the antibacterial activity of both the crude and purified Methanolic extracts of *P. murex* fruits. For the antifungal assay, PDA medium was prepared supplemented with five concentrations (0.01-1.0 mg/ml)of the Methanolic extracts. Afterwards plates were inoculated with the plant pathogenic fungi as3 mm^2^ plugs placed at equal distances (2-2.5 cm). Radial growth was measured after the 3^rd^ day of incubation of 24 h in BOD at 25°C. Control had six inoculations of phytopathogen on a plate supplemented with solvent only. Growth Inhibition (GI%) was calculated using the following formula: GI= [(A-B)/A]x 100 Where A= diameter of fungi in the solvent only; B= diameter of fungi in fruit extract intoxicated plates at various concentrations. Inhibitory concentration (IC_50_) was calculated by regression equation analysis (Chaudhary and Kaushik, 2015).

For antibacterial assays, 22.5 ml of NAM was poured on sterile Petri dishes (90 mm) and allowed to solidify. Agar surface of each plate was streaked by a sterile cotton swab with the reference bacterial strain. A paper disc of 6 mm size was impregnated in 100 μL of each sample and was placed on solidified agar plates at equal distance with control. The plates were allowed to stand for 30 min. After that, the plates were incubated at 28°C for 24 h for the growth of bacteria.

### Statistical analysis

All the results were expressed as the average ± Standard Error of six replicates. An one-way ANOVA was performed to check the significance of the results obtained at p-value ≤0.5.

### TLC and HPLC analysis

Isolation and purification of the compound from active extract was done using preparative TLC. Approximately, 200 mg of Methanolic crude extract was loaded on the Merck TLC Silica gel 60 (20 × 20 cm) plates and run in solvent system Ethyl Acetate: Hexane (6:4) in a TLC chamber. Ultraviolet light (365 and 366 nm) was used for the screening of those bands which were not visible in day light on TLC plates. For each band, Rf value was calculated, and the band was scrapped individually. Each collected bands from TLC plates were washed thrice with the HPLC grade Methanol (Fisher scientific, Qualigens) and concentrated in a rotory evaporator (Heidolph, Germany).

### HPLC analysis

HPLC of the methanolic extracts and of all the collected TLC fractions was performed on HPLC system equipped with Autosampler 717 plus; PDA 2996 detector and Empower2 software (Waters, USA). The samples were analyzed on an RP-18 column (250×4.60 mm, five μ), using Acetonitrile and HPLC grade water gradient as mobile phase (Chaudhary and Kaushik, 2015). The sample for HPLC was prepared by dissolving 1mg of the extract in 1ml of HPLC grade methanol.

### Spectroscopic analysis of the isolated compound

Approximately, 5-6 mg of the compound isolated by HPLC was used for the characterization by FT-IR and NMR, analysis.

### FTIR Spectrum analysis

The FT-IR spectrum was obtained using a Nicolet 380 (Thermo Scientific, USA) spectrometer at room temperature. The functional group was identified using potassium bromide (KBr) pellets and scanned in the range of 4000-400 cm^−1^.

### ^13^C NMR and ^1^H NMR analysis

Approximately 6 mg of samples were dissolved in 600 μl DMSO, vortexes (1 min) and sonicated (5 min), After sonication sample was vortexed (1 min) again and centrifuged (13,200 rpm). The supernatent was transferred to a 5mm NMR tube. ^1^H NMR spectra was referenced internally on CDCl_3_ (δ1 H 7,26) and ^13^C NMR spectra (Bruker spectrometer, Germany) was referenced internally on CDCl_3_ (δ13C 77.20). Chemical shifts (*δ*) are expressed in ppm, with the coupling constants (*J*) reported in Hertz (Hz).

### LC/MS analysis

LC-MS experiments were performed on a liquid chromatographic/autosampler system that consisted of a Waters Alliance 2690 Separations Module (Waters, Milford, MA) and a Micromass LC-TOF™ II mass spectrometer (Micromass, Wythenshawe, UK) equipped with an orthogonal electrospray source (Z-spray).

### The optical activity

Optical rotation was determined on a Autopol V Polarimeter. The substance was stored in a 0.5 ml cuvette with 0.1 dm length the angle of rotation was measured at a wavelength of 546 and 579 nm of a mercury vapour lamp at room temperature (25°C). The optical rotation at the wavelength 579 and 546 nm respectively, calculated using the formula:

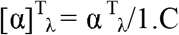

Where: *α* = the measured angle of the rotation in degrees I = the length in dm of the polarimeter tube, C = the concentration of the substance expressed in g/100 ml of the solution.

## Results

### Screening of antifungal and antibacterial activity of different fruit extracts

Antimicrobial activity of 29 fruits extracts of *P. murex* collected from different locations of India (Table S1) was screened against eight plant pathogenic fungi and four plant pathogenic bacteria (Table S2). Crude extracts of fruits were isolated in Methanol (Table S3) and diluted to specific range of concentrations (0.001 to 1mg/ml) to perform the bioassay.

Among the 29 fruits extracts assayed against 12 microbes, only 11 fruit extracts were found to contain some bioactive molecules and that too only against three microbes viz. *R. solani, B. cinerea*, and *S. sclerotiorum* (Table 1 and Figure 1). Among the 11 samples, Asakhedi village and Indore smaple showed the highest GI (%) i.e. ~23.21 and ~25, respectively while Neemunch showed least GI (%) at lowest concentrations (10μg/ml) of the crude fruit extracts. Till 100 μg/ml concentrations wide variations in GI (%) has been observed which become more stable at higher concentrations i.e. at 500 or 1000 μg/ml. According to GI (%) and IC50 value, five extracts were found positive against *R. solani*, and seven extracts were found active against *S. sclerotiorum* while only one extract was found active against *B. cinerea* (Table 1 and Figure 1).

**Figure 1.**
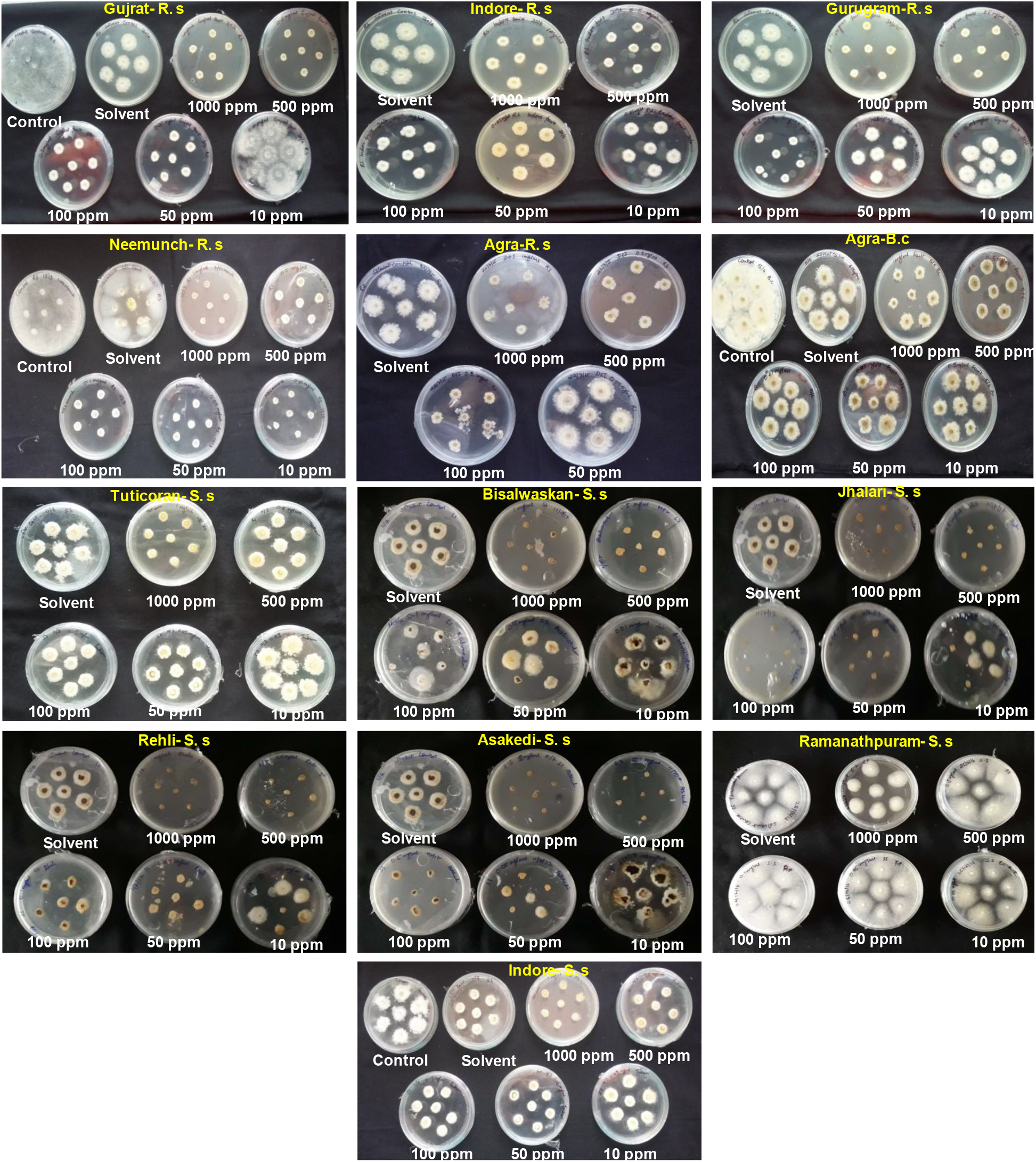
Fungal growth assay showing inhibitory effects of methanolic friut extracts isolated from different regions on three fungus R.s. (*Rhizoctonia solani*), B.c. (*Botrytis cineara*) amd S.s. (*Scleratonmnium sclerotium*) at various concetartions of crude extracts.

**Table 1.**
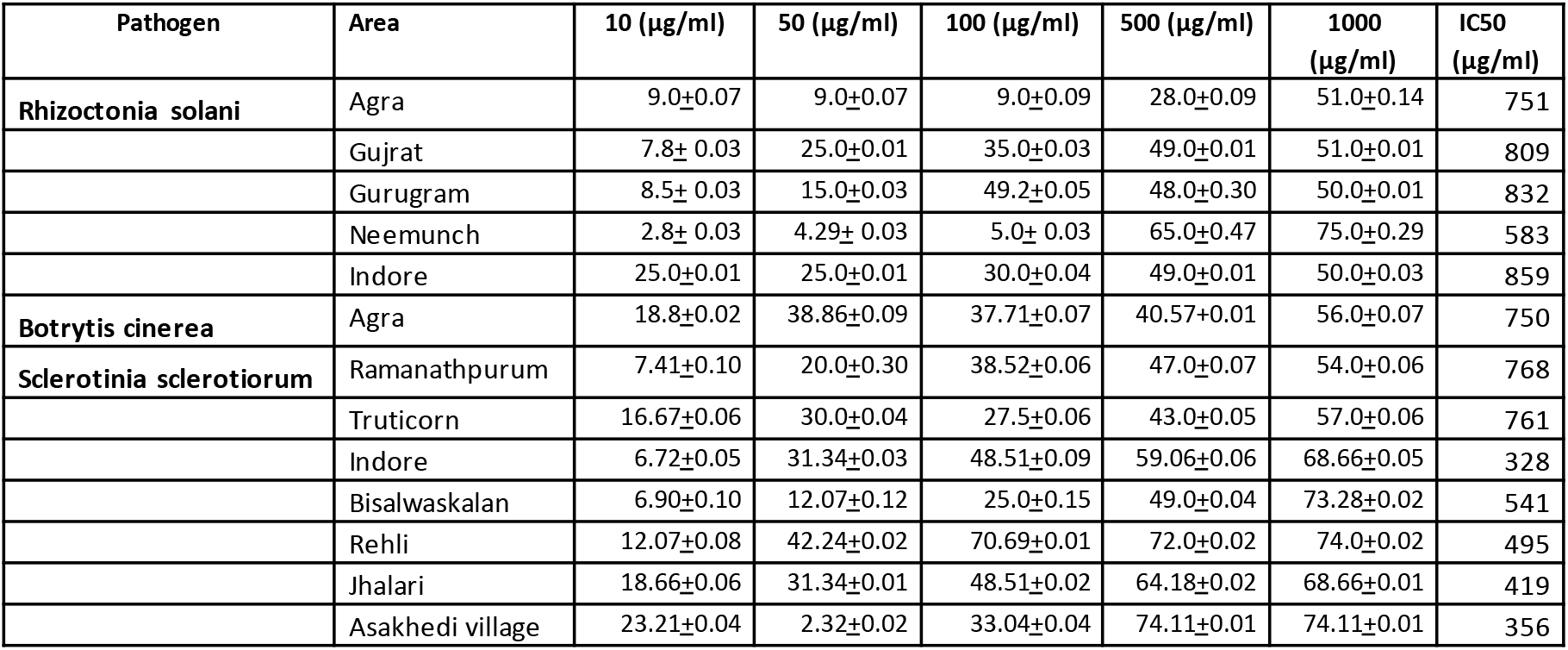
GI (%) and IC50 values obtained from the bioactivity assay of crude fruit extract on fungal species

| Pathogen | Area | 10 (µg/ml) | 50 (µg/ml) | 100 (µg/ml) | 500 (µg/ml) | 1000 (µg/ml) | IC50 (µg/ml) |
| --- | --- | --- | --- | --- | --- | --- | --- |
| **Rhizoctonia solani** | Agra | 9.0±0.07 | 9.0±0.07 | 9.0±0.09 | 28.0±0.09 | 51.0±0.14 | 751 |
|  | Gujrat | 7.8± 0.03 | 25.0±0.01 | 35.0±0.03 | 49.0±0.01 | 51.0±0.01 | 809 |
|  | Gurugram | 8.5± 0.03 | 15.0±0.03 | 49.2±0.05 | 48.0±0.30 | 50.0±0.01 | 832 |
|  | Neemunch | 2.8± 0.03 | 4.29± 0.03 | 5.0± 0.03 | 65.0±0.47 | 75.0±0.29 | 583 |
|  | Indore | 25.0±0.01 | 25.0±0.01 | 30.0±0.04 | 49.0±0.01 | 50.0±0.03 | 859 |
| **Botrytis cinerea** | Agra | 18.8±0.02 | 38.86±0.09 | 37.71±0.07 | 40.57±0.01 | 56.0±0.07 | 750 |
| **Sclerotinia sclerotiorum** | Ramanathpurum | 7.41±0.10 | 20.0±0.30 | 38.52±0.06 | 47.0±0.07 | 54.0±0.06 | 768 |
|  | Truticorn | 16.67±0.06 | 30.0±0.04 | 27.5±0.06 | 43.0±0.05 | 57.0±0.06 | 761 |
|  | Indore | 6.72±0.05 | 31.34±0.03 | 48.51±0.09 | 59.06±0.06 | 68.66±0.05 | 328 |
|  | Bisalwaskalan | 6.90±0.10 | 12.07±0.12 | 25.0±0.15 | 49.0±0.04 | 73.28±0.02 | 541 |
|  | Rehli | 12.07±0.08 | 42.24±0.02 | 70.69±0.01 | 72.0±0.02 | 74.0±0.02 | 495 |
|  | Jhalari | 18.66±0.06 | 31.34±0.01 | 48.51±0.02 | 64.18±0.02 | 68.66±0.01 | 419 |
|  | Asakhedi village | 23.21±0.04 | 2.32±0.02 | 33.04±0.04 | 74.11±0.01 | 74.11±0.01 | 356 |

Other 9 microbial species does not showed any alteration in the growth pattern hence not presented here (data not shown). Moreover, interesting observation is that fruit extract samples from one location respond to specific fungi, only and remains ineffective for other fungi. Contrary to it, Agra and Indore sample showed activity against two different fungal species. Like, the extract from Indore location was active against both *R. solani* as well as against *S. sclerotiorum*. Hence, considering activity against broad spectrum fungi we selected sample from Indore region for downstream analysis for compound characterization.

### Isolation and purification of the bioactive compound

Because of broad spectrum action of fruit extract sample from Indore we selected it for further downstream extraction of the biochemical molecule and its structural elucidation. For the compound isolation, the extract was fractionated by preparative TLC. A total of Nine different bands were detected on the TLC plate under visible and UV light, with Rf value corresponding to 0.029, 0.076, 0.194, 0.235, 0.318, 0.441, 0.482, 0.558, 0.91(Fig. S1). All these bands were scrapped and extracted in Methanol for analysis using HPLC to detect presence of either single or multiple compounds (Fig. 2). Among the Nine samples analyzed, we detected presence of a sharp and single peak in band that corresponds to Rf value 0.318 in TLC analysis (Figure 2). Other fractions from TLC plates yielded mixture of peaks revealing presence of multiple and complex molecules (Fig. 2).

**Figure 2.**
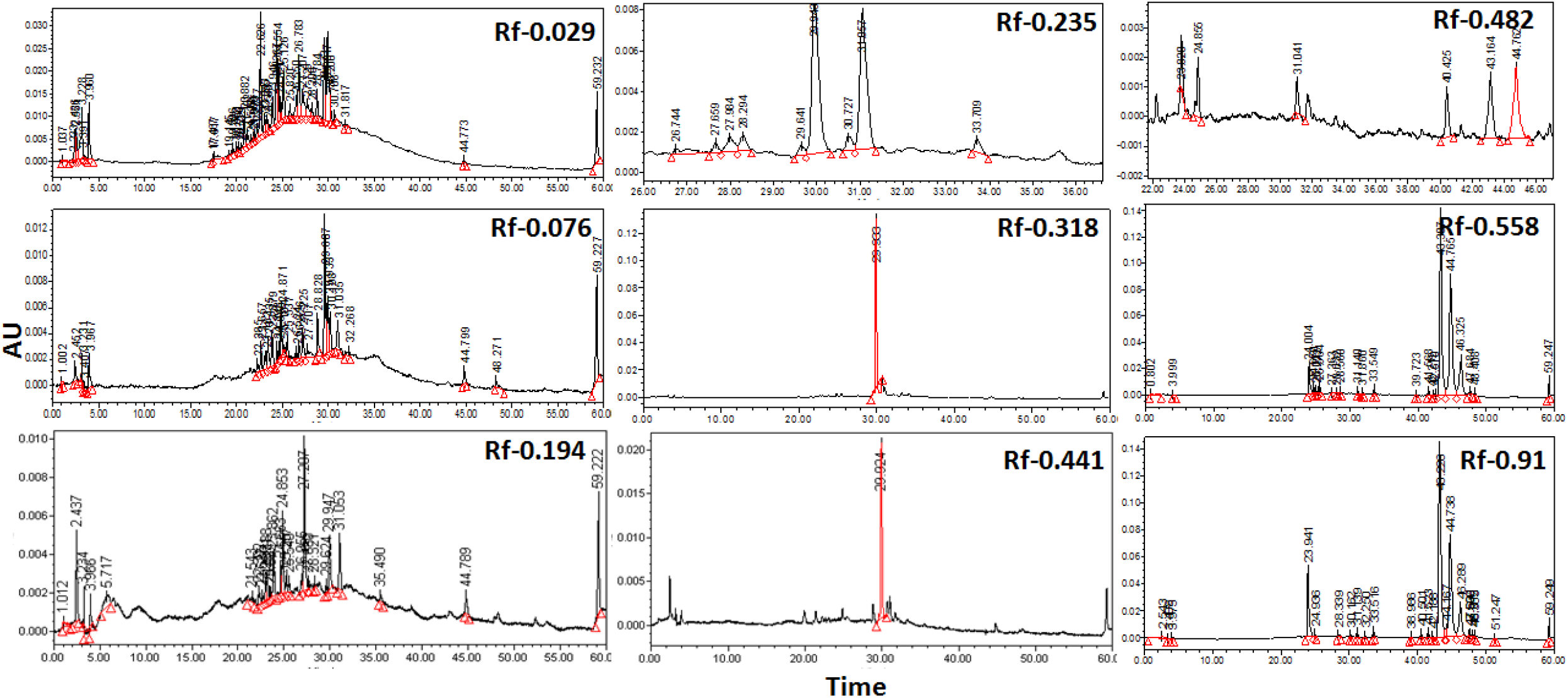
HPLC chromatograms of crude methanolic extracts of different bands appeared on TLC plates corresponding to different Rf values.

### Bioactivity estimation of the fractionated compound

Before structural elucidation, we evaluated different all Nine TLC Methanolic fractions for presence of bioactive compounds through bioassay against the already screened microbes i.e. *R. solani, B. cinerea*, and *S. scleratonium*. Among all the samples analyzed band corresponding to Rf value 0.318 showed significant potent activity against the fungal phyto-pathogen viz. *S. scleratonium* having IC_50_ value at 454 μg/ml (Figure 3; Table 2) by suppressing its mycelia at 5.7 mm of growth radius. Other species (*R. solani* and *B. cinerea*) which were found susceptible to the crude extracts were not found susceptible against this particular fraction of metabolites.

**Figure 3.**
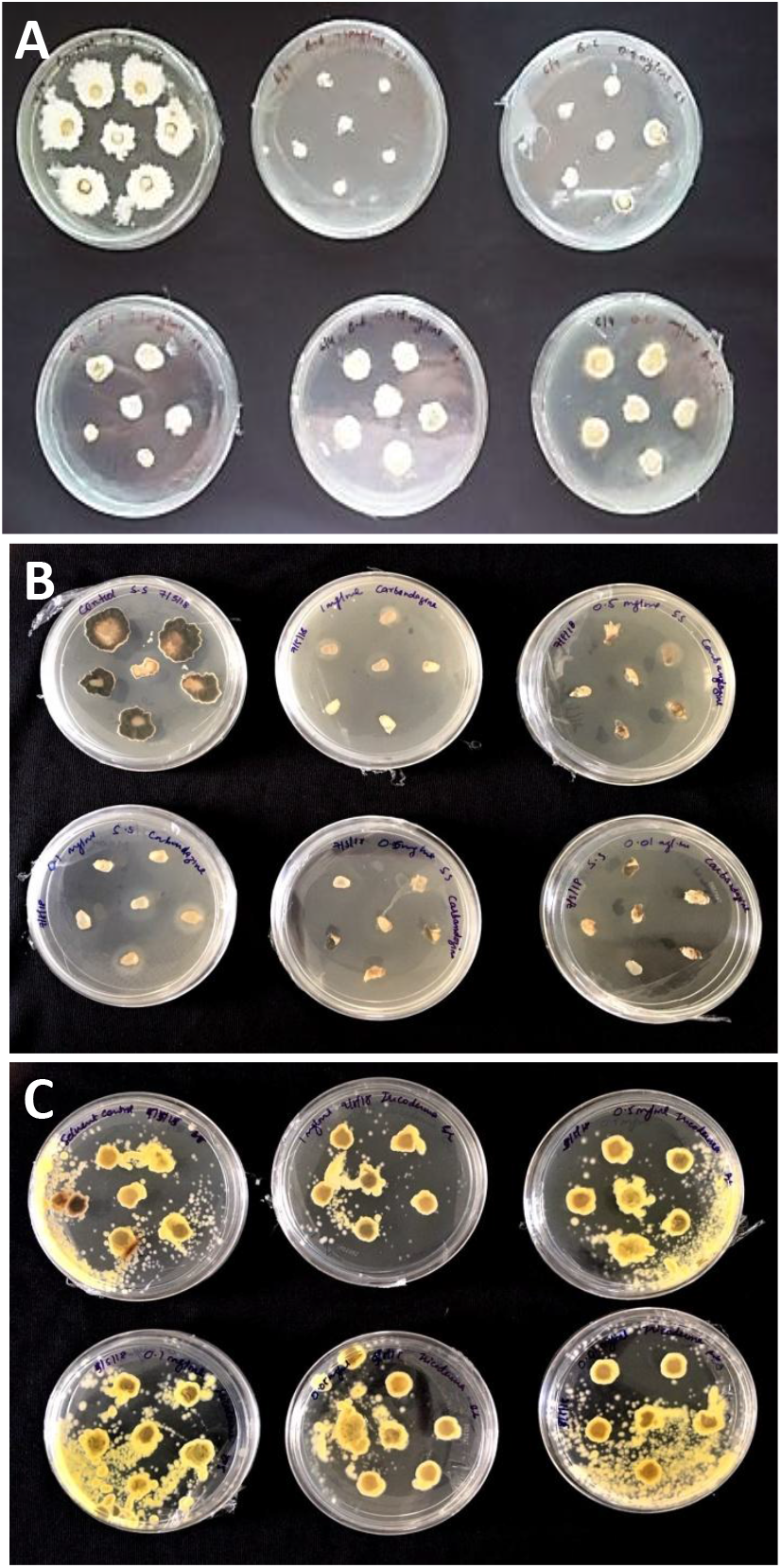
Fungal growth assay of A) *S. scletonia* on medium supplemented with purified HPLC fraction corresponding to 5th TLC fraction with Rf value 0.318; B) *S. scletonia* on medium supplemented with Carbendazim at various concetartions; C) *Trichoderma* sp. showing normal growth on HPLC purified extract.

**Table 2.** GI (%) values obtained from the bioactivity assay of purified compound on *S. sclerotiorim*

| Pathogen | Concentration (µg/ml) | Growth inhibition (mm) ±SE |
| --- | --- | --- |
| *Sclerotinia sclerotiorum* | 10 | 18.3±0.03 |
|  | 50 | 12.7±0.02 |
|  | 100 | 11.9±0.01 |
|  | 500 | 7.9±0.03 |
|  | 1000 | 5.7±0.03 |

To ensure the quality of our assay we analyzed another potent fungicide Carbendazim, which is a widely used, broad-spectrum benzimidazole fungicide under similar conditions and found it highly potent against the microbes as it limits the growth at lowest concentrations. Contrary to it, we checked for effect of isolated compound on beneficial fungi like *Trichoderma*. At different concentration of the isolated compound we could not found any significant difference in the growth pattern of the *Trichoderma* species.

### Mass estimation of the bioactive compound

The isolated fraction was further characterized by the FTIR analysis. Data of IR spectrum signals were determined for several groups; at 3015 representing CH; at 2945, 2837, 1735, 1641 representing C=O; and at 1510, 1427, 1415, 1085 representing C-O respectively (Fig. 4).

**Figure 4.**
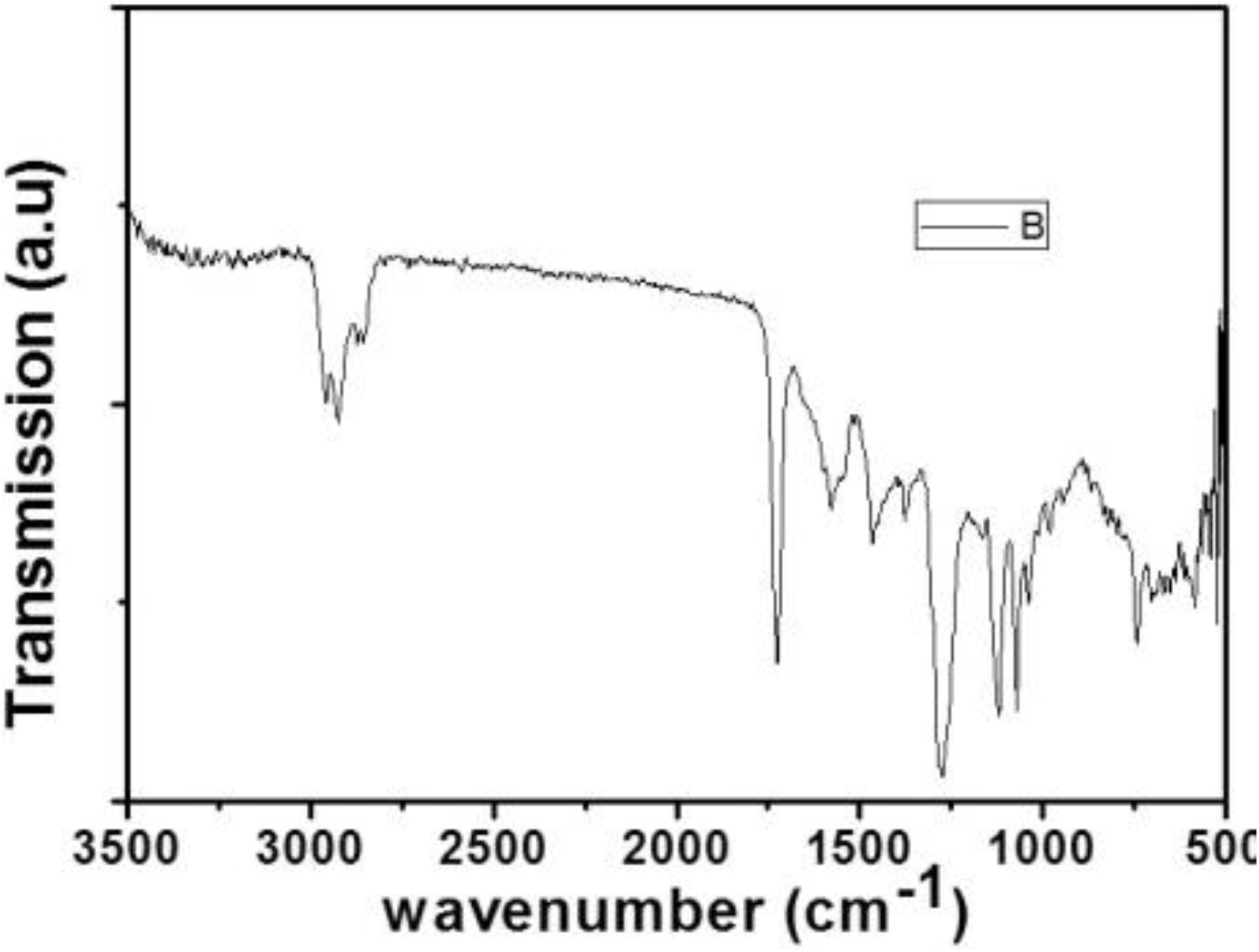
FTIR spectrum of the HPLC purified isolated compound

### Structural Elucidation of the compound

In the proton spectrum, the proton of C2 appeared as at triplet peak at δ 4.09 with j=6.0Hz, while C3 shows triplet peak at 0.99 with J-=6.5 Hz and methanol at 10’ At 1 C shows the broad multi-peak at j=18.8Hz. C7 shows C=O at 167.72 (Figure 5A). ^13^C NMR of the compound showed carbon resonance at δ 167.72 by two bond which was further confirmed by IR the presence of carboxyl group, ketone, aromatics, and also reveals the other stretching and bending vibrations of functionalities in the compound (Figure 5B). Its mass spectrum showed the molecular ion peak at m/z 320 [M]+, which correspond to the molecular formula of compound (C_20_H_32_O_3_) (Figure 5C). The compound get fragmented at (14.5), 215 (2.7), 207 (10.3), 199 (3.2), 177 (3.6), 143 (45.1), 121 (2.8), 113 (59.2), 105 (73.5) (Figure SA&B). On the basis of these observations, the purified compound is purposed to be 5- β -Hydroxyltridecayl benzoate (Figure 5C) with at optical rotation 0.143°.

**Figure 5.**
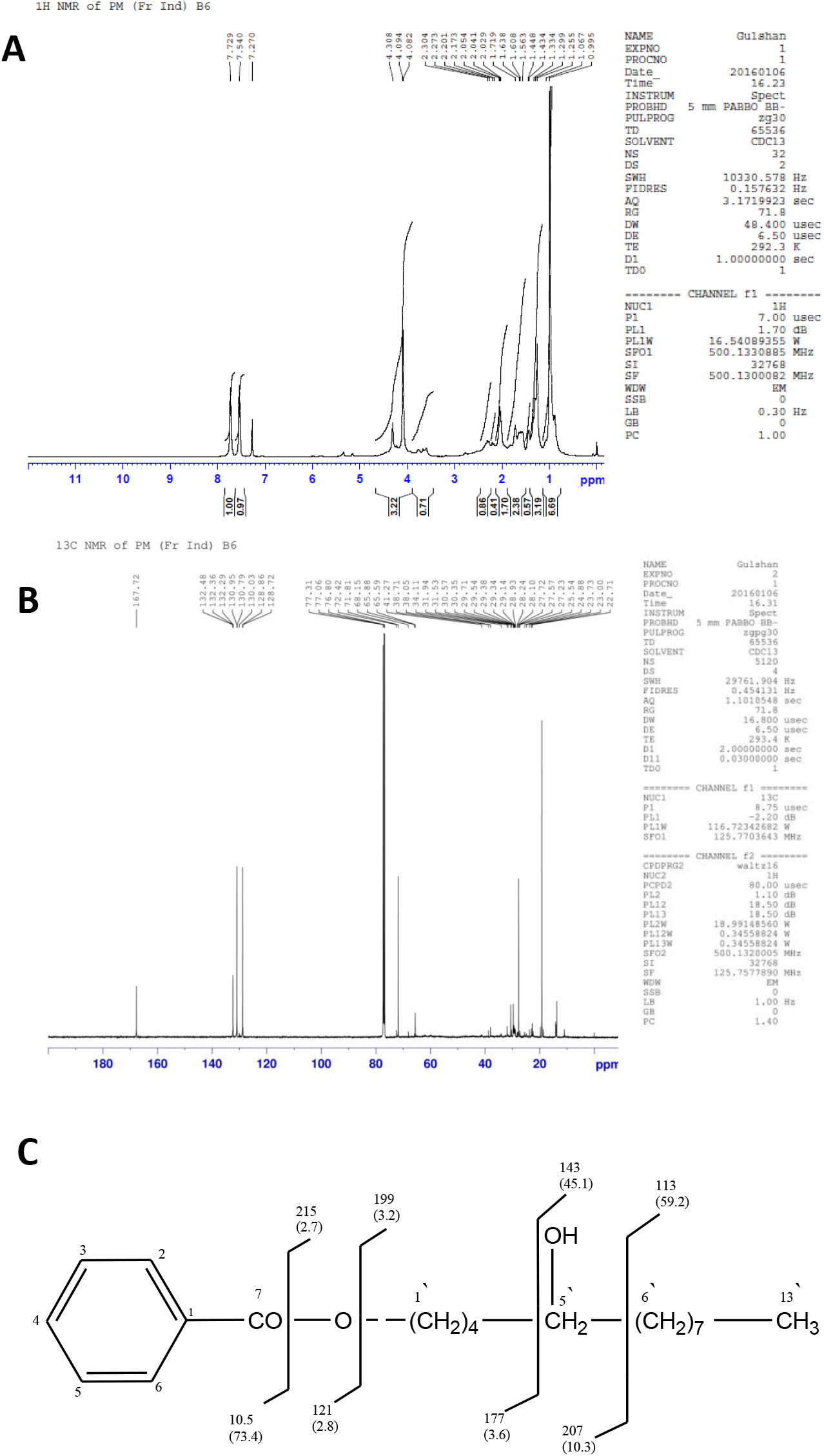
Structural analysis of the compound using NMR. A) proton NMR; B) Carbon NMR; C) Structure and fragmentation analysis of the deduced compound.

## Discussion

The use of synthetic fungicides has raised concerns about environmental issues. In this study, the methanolic fruits crude extract of *P. murex* showed potent activity against the *R. solani, B. cinerea* and *S. sclerotiorum*. The impacts of these phytopathogens are severe to influence the quality and yield of the crops (Yazdani et al., 2011).

The crude sap, volatile and essential oil extracted from whole plant or specialized plant parts are widely used in preparing the antimicrobial compounds which are significantly used against the different plant pathogens/diseases. *P*lants can synthesize aromatic secondary metabolites, most of which are phenols or their oxygen-substituted derivatives namely, quinones, flavones, flavonoids, flavonols, tannins and coumarins (Saravanakumar et al. 2015). These groups of compounds exhibits antimicrobial effect because of number of hydroxyl or carbonyl group (-OH or C=O) groups present in the compound. In general, the longer chain (C6–C10) molecules in plant extracts have been observed to have greater antifungal properties (Ultee et al. 2002; Holley and Patel 2005). They are synthesized by plants in response to microbial infection and are often found effective *in vitro* as antimicrobial substance against a wide array of microorganisms.

In the present study, the crude extract from *P. murex* fruits has lower IC_50_ value than the isolated compound. The isolated compound from the Indore location strongly inhibits particular plant fungi *S. sclerotiorum* than other Eleven examined plant pathogens. From the published data, it was found that there are some bioactive compounds which are active against several phytopathogens such as *R. solani, B. cinerea, S. sclerotiorum* because the bioactive compounds have the presence of some functional groups such as Phenolics, Tannins, flavonoids, etc.(Rai and Carpinella 2006). The mechanism of action of plant products on fungal cells is thought to be: (i) granulation of cytoplasm; (ii) membrane rupture in cytoplasm; (iii) inhibition and inactivation of intracellular and extracellular enzyme synthesis. These actions can occur in an isolated or a concomitant manner and culminate with mycelium germination inhibition (Cowan1999).

Earlier bioactive compounds like tannins from *L. tridentate* and *F. cernua* (Castillo et al. 2010); Daturilin extracted from the leaves of *Datura metel* (Kagale et al., 2011) is reported to be active against pathogen *R. solani*. Another bioactive compound *eicosane* and dibutyl phthalate isolated from microbial species *Streptomyces* strain KX852460 reported as a potential bio-control agent for *R. solani* (Ahsan et al. 2017).

The compound eugenol has reported for the direct inhibition growth of *B. cinerea in vitro* (Wang et al. 2010). The essential oil from the leaves of *Ocimum sanctum* Linn., *Prunus persica* Linn is reported to be active against *B. cinerea* of grapes (Tripathi et al. 2008). The Essential oils Limonene, Linalool. Myrcene isolated from epicarp of *Citrus sinensis* is reported to be active against *Botrytis cinerea* in tomato, mango (Sharma and Tripathi 2006). Polyphenols and flavonoids from Methanolic leaf extract of Zizyphus is active against *Botrytis cinerea* (El-Khateeb et al., 2013). A novel anti-phytopathogenic compound 2-heptyl-5-hexylfuran-3-carboxylic acid was isolated from the fermentation broth of *Pseudomonas* sp. SJT25 is reported active against phytopathogenic *B. cinerea* and *R. solani* (Wang et al., 2011). Minocycline was considered as the primary active molecule produced by ENT-18 against *S. sclerotiorum* (Zucchi et al., 2014). In present studies targeted extract from Indore location, a novel compound isolated and characterized by using several techniques as 13C NMR, 1H NMR, FTIR, and named 5- β - Hydroxyltridecayl benzoate. This isolated compound is a novel compound as it is first time derived and reported from the extract of *Pedalium murex* fruits. After isolation this compound is screened against *S. sclerotiorum B. cinerea* and *R. solani* and found effective against *Sclerotinia sclerotiorum*, pathogenic fungi cause white mold disease. The characterized structure derived from the extract having the benzene ring with CH group this may be responsible for the granulation or membrane rupture in the cytoplasm; or inactivation of intracellular and extracellular enzyme synthesis that resulted in mycelium germination inhibition.

Recently, cuminic acid isolated from the seed of *Cuminum cyminum* L is reported for its activity against *S. sclerotiorum as* it exhibited significant control effects against *S. sclerotiorum* (Sun et al., 2017). Botanical fungicide cuminic acid as p-isopropyl benzoic acid is a natural alternative to commercial fungicides or a lead compound for the control of *Sclerotinia* stem rot (Sun et al., 2017; Huzar-Novakowiski et al. 2017). While According to Goussous et al. 2013 the extracts of rosemary and Greek sage leaves are highly active against *S. sclerotiorum* and states that it could become natural alternatives to synthetic fungicides to manage diseases of *S. sclerotiorum*. The crude extract from Indore is active against two phytopathogens *R. solani* and *S. sclerotiorum* with a low IC_50_ value less than the isolated compound. To identify another compound responsible for the inhibition of *R. solani* need to know in further research. The suppression of phytopathogen might be due to the loss of cell membrane integrity or breakage of cell membrane due to the release of certain chemicals (Giorgio et al. in 2015). The present study first time reveals the potential of isolated compound 5- β -Hydroxyltridecayl benzoate from *P. murex* fruits extract against phytopathogen, *S. sclerotiorum*. This study needs more research in this field to carry out *in vivo* experiments because in future it can be exploited as bio fungicides for the agriculture sector as this study paves the way for the development of bioactive natural products/fungicide with the added benefits of an environmentally safe and economically viable product

## Acknowledgement

Authors thankful to Dr. M. Ali for help in data analyzing AIRF lab for NMR, LCMS facility, IIT department of chemistry for providing polarimeter and FTIR facility, New Delhi for providing The authors are also thankful to Director General, TERI, for providing facilities.

## Funding Agency

UGC Women’s Post Doctorate Fellowship GE-UTD-6589.

## Compliance with Ethical Standards

### Conflict of interest

All authors has no conflict of interest

### Ethical approval

This article doesn’t contain any studies with human participants performed by any of the authors.

## Glossary

GI: Growth Inhibition
IC_50_: Inhibition concentration
PDA: Potato Dextrose Agar media
NAM: Nutrient Agar Media
TLC: Thin Layer Chromatography

